# Tagger: BeCalm API for rapid named entity recognition

**DOI:** 10.1101/115022

**Authors:** Lars Juhl Jensen

## Abstract

Most BioCreative tasks to date have focused on assessing the quality of text-mining annotations in terms of precision of recall. Interoperability, speed, and stability are, however, other important factors to consider for practical applications of text mining. The new BioCreative/BeCalm TIPS task focuses purely on these. To participate in this task, I implemented a BeCalm API within the real-time tagging server also used by the Reflect and EXTRACT tools. In addition to retrieval of patent abstracts, PubMed abstracts, and PubMed Central open-access articles as required in the TIPS task, the BeCalm API implementation facilitates retrieval of documents from other sources specified as custom request parameters. As in earlier tests, the tagger proved to be both highly efficient and stable, being able to consistently process requests of 5000 abstracts in less than half a minute including retrieval of the document text.

## 1 Introduction

BioCreative and other shared tasks in the biomedical text-mining community have over the years played a key role in progressively improving text-mining methods, in particular for named entity recognition (NER). Most BioCreative tasks have focused purely on evaluating the precision and recall (1,2), with the BioC interoperability task (3) and the interactive annotation task (IAT) (4) being notable exceptions. However, as illustrated by the latter two tasks, whereas precision and recall are obviously important factors, they are far from the only factors that matter when using text mining in practice. Interoperability, speed, and stability are other very important factors; the new Technical Interoperability and Performance of annotation Servers (TIPS) task set out to evaluate just that.

I participated in the BioCreative V IAT (4) with the interactive annotation tool, EXTRACT, which helps curators find and extract standard-compliant terms for annotation of metagenomic records and other samples (5). Behind its web-based user interface, the system makes use of the same real-time tagger for NER as the augmented browsing tool Reflect (6). The core NER engine was designed from the ground up with speed in mind and is capable of tagging thousands of PubMed abstracts per second per CPU core (7). This makes it ideally suited for large-scale and real-time applications, such as the TIPS task.

Here, I present a BeCalm API for the NER tagger underlying the EXTRACT (5) and Reflect (6). The system delivered a total turnaround time of about 1 second for small requests, and was able to process approximately 5,000–10,000 abstracts per minutes for larger batch requests. Notably, the vast majority of this time was spent on retrieving the document text rather than actual processing of it; to make the server faster, it would thus be necessary to locally cache the documents, which was explicitly not permitted in the TIPS task.

## 2 Materials and Methods

### Dictionaries used for NER and normalization

The server uses a combination of previously published dictionaries to recognize six of the types of entities accepted by the BeCalm server and normalize them to identifiers from databases and ontologies. These are a subset of the entity types used in EXTRACT v2 (5).

For annotation of gene/protein names, the tagger uses a dictionary covering the 9.6 million protein-coding genes from 2031 0.organisms included in STRING v10.5 (8) as well as ncRNAs from the RAIN database (9). Unlike many NER systems, the BeCalm API makes a distinction between genes and their protein products. Because the STRING database is locus-based, i.e. it does not distinguish between splice isoforms, and because ncRNAs are also included, I chose to use the type GENE for these annotations and to not support the PROTEIN annotation type. All recognized names are disambiguated to their respective STRING or RAIN identifiers, which are derived from the Ensembl (10), RefSeq (11), and miRBase (12) databases.

Annotations of the type CHEMICAL are made using a dictionary comprised of small-molecule compounds from the PubChem database (13), which was developed and used for recognition of chemical names in STITCH v5 (14). All annotations of chemicals are normalized to PubChem compound identifiers.

The tagger makes annotations of the type ORGANISM using an updated version of the dictionary of the SPECIES/ORGANISMS tagger (7). The dictionary was constructed based on NCBI Taxonomy (10), and all annotations are thus normalized to NCBI taxon identifiers.

For SUBCELLULAR_STRUCTURE, TISSUE_AND_ORGAN, and DISEASE the tagger uses the dictionaries created as part of the COMPARTMENTS (15), TISSUES (16), and DISEASES (17) database, respectively. These were constructed from Gene Ontology (18), Brenda Tissue Ontology (19), and Disease Ontology (20), identifiers from which are used for normalization of the annotations.

The version of the dictionary used by Tagger for the TIPS task has been deposited on FigShare (doi:10.6084/m9.figshare.4578292). The reduced dictionary used by PiTagger has also been deposited on Figshare (doi:10.6084/m9.figshare.4635175). The latest version of the dictionary, which is used by the production server, is available for download at http://download.jensenlab.org/tagger_dictionary.tar.gz.

### Named entity recognition software

The core of the NER system is a highly optimized dictionary-based tagger engine, implemented in C++. It is able to perform flexible matching of a dictionary with millions of names against thousands of abstracts per second per CPU core (7). The tagger is furthermore inherently thread safe, for which reason a single instance of the tagger can easily handle many parallel requests. These properties make it an excellent starting point for building a real-time service that can handle large requests as required for TIPS task.

Although the TIPS task does not assess the quality of the annotations, it is worth nothing that the speed of the tagger was not achieved by sacrificing quality. The quality of the tagging results for organism names was previously evaluated on gold-standard corpora and found to be comparable to the best methods (7,21). The NER quality has not been benchmarked directly for chemicals, genes, tissues, and diseases has not been benchmarked directly; however, these NER components have shown to give good results when used as the basis for association extraction (8,9,13,15–17).

The tagger software is open source and available at https://bitbucket.org/larsjuhljensen/tagger/. It can be used either as a command-line tool or as a Python module. It is also distributed as a Docker container at https://hub.docker.com/r/larsjuhljensen/tagger/.

### BeCalm API implementation and hosting

I implemented the BeCalm API itself in Python and runs as a module under an in-house web service framework. The framework uses multiple queues and thread pools to simultaneously run several compute intensive requests in in parallel (e.g. getAnnotations requests) and be responsive to smaller requests (e.g. getStatus). The API code accesses a single instance of the tagger engine through its Python module, which has the complete dictionary preloaded in memory.

The main tagger runs on a single server with one Intel Xeon E5-2620 2.4 GHz CPU and 256 GB of RAM. This server also runs many other resources and databases related to text mining, including EXTRACT (5), SPECIES/ORGANISMS (7), COMPARTMENTS (15), TISSUES (16), and DISEASES (17). This server is physically hosted at the high-performance computing facility Computerome and is from hereon referred to as Tagger.

To test the influence of the performance of actual document tagging vs. overhead associated with fetching of document texts, I ran a second instance of the tagger on a Raspberry Pi 3 with a 1.2 GHz quad-core ARM Cortex-A53 and 1 GB and RAM. Due to the limited memory, this instance runs with a reduced dictionary; however, it should be noted that tagging speed is largely independent of dictionary size because the tagging algorithm is based on hash lookups (7). This instance was hosted over my home internet connection (60 Mbit/s download, 25 Mbit/s upload) and is in the following referred to as PiTagger.

## 3 Results and Discussion

### Rapid annotation of biomedical entities

To test the speed of Tagger and PiTagger when accessed through the BeCalm API, I submitted private requests for tagging of 1, 10, 100, 1000, and 5000 abstracts from the abstract and patent servers via the BeCalm web interface.

All settings except from the number of documents to tag were left at their default values. Each of the five sizes of tagging requests was repeated five times at four different timepoints, giving a total of 20 observations of the total time required for tagging for each size of request from each document source on each of the two tagger servers. These results are summarized as means and standard deviations in Table 1.

**Table 1.**
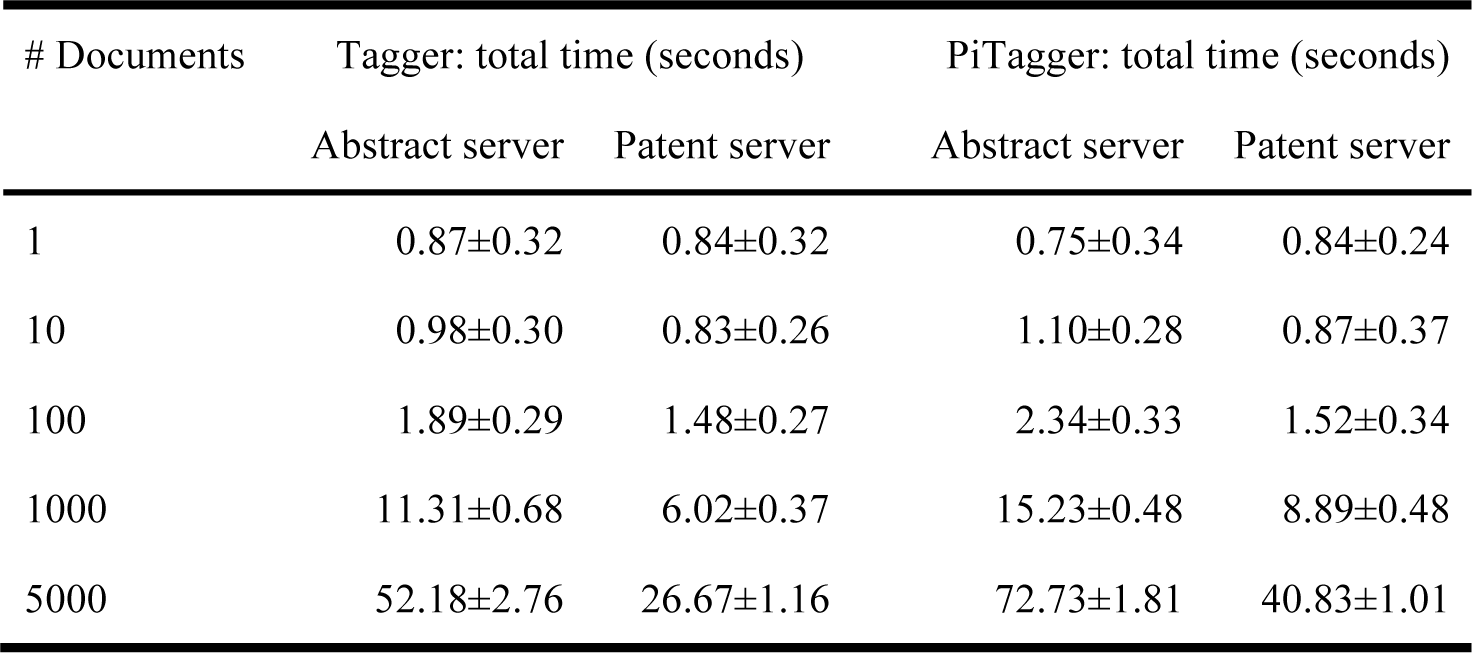
Performance of the taggers. For small requests the total turnaround time is ~1 second. Larger requests take an extra 5–10 seconds per 1000 abstracts to be processed on Tagger. Notably, most of this time is spent on retrieving the document texts from document identifiers, whereas the actual NER step takes only about 20% of the total time. This is reflected in the fact that the PiTagger, which runs on a Raspberry Pi 3, takes only about 50% longer to process large requests.

Neither Tagger nor PiTagger suffered any errors or slowdowns during these tests, despite the Tagger server hosting multiple other resources and the PiTagger running on minimal hardware. This shows that the software is not only fast but also stable. This is unsurprising since all parts except the BeCalm API-specific code have been used in a production setting for several years.

In summary, the Tagger speed tests showed that there is a constant overhead of about 1 second on all tagging requests, which dominates the picture up to tagging of about 100 patent abstracts. For larger requests, the service takes ~5 and ~10 seconds more per 1000 patent abstracts and PubMed abstracts, respectively. This difference is presumably explained by PubMed abstracts being, on average, about twice as long as patent abstracts. Notably, the vast majority of the time is spent on fetching the document texts, with only about ~20% of time being spent on actual processing. Although explicitly not permitted in the TIPS task, local storage or caching of documents on the annotation server would thus be an attractive future feature.

To further test and illustrate that retrieval of document texts is the main bottleneck, I configured a second copy of the tagger code, PiTagger, to run on a Raspberry Pi 3. For small requests, the total time is indistinguishable between Tagger and PiTagger, and even for large requests PiTagger takes only about 50% longer than Tagger (Table 1). This is the case despite the service running only one thread per request, thus utilizes only a quarter of the compute power of a Raspberry Pi 3 in these tests. The PiTagger did not participate in the full official TIPS evaluation.

The total tagging time for the official TIPS requests was in the beginning consistently longer than for the private requests reported in Table 1, which were submitted during the same weeks. Monitoring the tagging services during TIPS requests revealed that actual document processing was as fast as always. In light of the results above, I assume that this slowdown was due to the fetching of documents taking longer in the official tests, because all participants simultaneously send requests to the central document servers.

### Extending the BeCalm API

The BeCalm API in its current form has certain design constraints that limit from the flexibility and thereby usefulness of the annotation servers. Firstly, document text is not submitted as part of the request, but must instead be fetched from designated sources based on the submitted document identifiers. Secondly, the results cannot be returned directly to the end user, but must be returned to the central BeCalm server. Through creative use of the *custom_parameters* part of the request, I have circumvented both of these constraints.

Instead of hardwiring the annotation server to use only the abstract and patent servers provided by BeCalm, the relationships between *source* and server URL are specified within a *servers* subsection of *custom_parameters.* This enables end users to obtain the tagging results for any desired documents, provided they make the documents available through an API compatible with the one used by the BeCalm document servers.

Similarly, the annotation server is not hardwired to return the annotation results to the BeCalm server. Instead, the *saveAnnotations* request will be made to the URL specified in as *apiurl* in the *custom_parameters* section. This allows end users to set up their own server to receive the results directly, if they so wish.

## 4 Acknowledgment

This work was supported by the Novo Nordisk Foundation [NNF14CC0001]. Thanks to Helen V. Cook for improvements to the source code and documentation of the tagger, Sune Pletscher-Frankild for developing the Python framework under which the BeCalm API runs, and the organizers of the 3rd Biomedical Linked Annotation Hackathon (BLAH3), where the BeCalm API was developed.

